# Type-2 diabetes alters hippocampal neural oscillations and disrupts synchrony between hippocampus and cortex

**DOI:** 10.1101/2023.05.25.542288

**Authors:** Gratianne Rabiller, Zachary Ip, Shahram Zarrabian, Hongxia Zhang, Yoshimichi Sato, Azadeh Yazdan-Shahmorad, Jialing Liu

**Author notes:** **Corresponding author**: Dr. Jialing Liu, Department of Neurological Surgery (112C), University of California at San Francisco and Department of Veterans Affairs Medical Center, 1700 Owens Street, San Francisco, California 94158, USA.

## Abstract

Type 2 diabetes mellitus (T2DM) increases the risk of neurological diseases, yet how brain oscillations change as age and T2DM interact is not well characterized. To delineate the age and diabetic effect on neurophysiology, we recorded local field potentials with multichannel electrodes spanning the somatosensory cortex and hippocampus (HPC) under urethane anesthesia in diabetic and normoglycemic control mice, at 200 and 400 days of age. We analyzed the signal power of brain oscillations, brain state, sharp wave associate ripples (SPW-Rs), and functional connectivity between the cortex and HPC. We found that while both age and T2DM were correlated with a breakdown in long-range functional connectivity and reduced neurogenesis in the dentate gyrus and subventricular zone, T2DM further slowed brain oscillations and reduced theta-gamma coupling. Age and T2DM also prolonged the duration of SPW-Rs and increased gamma power during SPW-R phase. Our results have identified potential electrophysiological substrates of hippocampal changes associated with T2DM and age. The perturbed brain oscillation features and diminished neurogenesis may underlie T2DM-accelerated cognitive impairment.

## Introduction

The global prevalence of type 2 diabetes mellitus (T2DM) is on the rise (Kaiser, Zhang, & Der Pluijm, 2018), with over half a billion people already affected in 2018. T2DM is also a significant risk factor for a wide spectrum of neurological diseases including ischemic stroke, depression, and the Alzheimer’s disease (AD) (van Sloten, Sedaghat, Carnethon, Launer, & Stehouwer, 2020). T2DM accelerates normal brain aging by increasing gray matter atrophy 26% ± 14% faster than seen with normal aging, thus T2DM patients displayed a more rapid rate of cognitive decline than typically associated with natural aging (Antal et al., 2022). As such, T2DM potentiates the development of dementia (Yu et al., 2020).

Like age, diabetes has complex and profound effects on brain volume, neural activity, functional connectivity and cognitive function (Dorsemans et al., 2017; Zhang et al., 2015; Zhou et al., 2010). However, how T2DM and age interact to cause structural and network changes that may underlie cognitive dysfunction remains unclear. In humans, T2DM causes EEG rhythms to shift from higher to lower frequencies and reduces neural synchrony albeit to a lesser extent compared to other pathological cognitive aging disorders such as AD (Benwell et al., 2020; Dauwels et al., 2011). Experimental models using invasive recording modalities to probe deeper brain regions like the hippocampus (HPC) offer better opportunity to decipher the neurophysiological outcome of age and T2DM in learning and memory, circumventing the difficulty in assessing cognitive behavior in the T2DM models (Zakaria, Ahmad, & Qinna, 2021).

Recent evidence suggests that neurogenesis might compensate for the age-associated hyperexcitability of the CA3 area via a feed-forward inhibition mechanism (Berdugo-Vega et al., 2020), supported by evidence of direct contact between the filopodia of the mossy fiber terminals of the new neurons and parvalbumin interneurons (Leal & Yassa, 2015; Wilson, Gallagher, Eichenbaum, & Tanila, 2006). Increasing neurogenesis via overexpression of the cell cycle regulators CDk4/cyclinD1 not only rescued degraded navigational strategy and improved memory performance in the aged mice, but also restored the profile of hippocampal sharp wave associated ripples (SPW-Rs) (Berdugo-Vega et al., 2020), a CA1 specific brain oscillation known to underlie memory consolidation. These findings suggest that reduced neurogenesis has a causal role in altering hippocampal trisynaptic circuitry, resulting in perturbed brain oscillations. Yet the role of T2DM in accelerating age induced reduction in neurogenesis and the extent in disturbing brain oscillations has not been established.

To test the hypothesis that T2DM-induced brain atrophy may alter connectivity between cortex and brain regions crucial for memory, we recorded field potentials in the sensorimotor cortex and HPC in two age groups of diabetic and normoglycemic mice under urethane anesthesia. We found that both aging and T2DM disrupted functional connectivity between the cortex and HPC and led to increase in the duration of SPW-Rs, and gamma power during SPW-Rs. These changes were associated with reduced neurogenesis in the HPC. However, compared to the impact of aging, T2DM additionally caused increased slowing scores and reduced aperiodic spectral exponent in the HPC. These results suggest that T2DM has a unique impact on brain oscillations and functional connectivity, beyond the effects of aging alone. The combination of these changes may contribute to the accelerated cognitive decline seen in older individuals with T2DM.

## Materials & Methods

### Animals and housing

Diabetic db/db mice (B6.BKS(D)-Lepr<db/db>/J) homozygous for the leptin receptor gene mutation were used as the model of obesity-induced T2DM, while heterozygous db/+ mice (B6.BKS(D)- Lepr<db/+>/J) were used as normoglycemic controls (Akamatsu et al., 2015; Kanoke et al., 2020; Nishijima et al., 2016). Male and female db/+ and db/db mice at 200 or 400 days of age were housed in the institutional standard cages (5 mice per cage) on a 12-h light/12-h dark cycle, with ad libitum access to water and food. All animal experiments were conducted in accordance with the Guide for Care and Use of Laboratory Animals issued by the National Institutes of Health and approved by San Francisco Veterans Affairs Medical Center Institutional Animal Care and Use Committee. The identity of each mouse subject was blinded to investigators who conducted the experiments and data analysis.

### Electrophysiological recording

Recordings were performed using 16-channel extracellular silicon electrodes (A1×16-5mm-100- 703, NeuroNexus Technologies) under urethane anesthesia (Sigma, 1 g/kg i.p.) for one hour (He et al., 2020; Ip et al., 2021). Following craniotomy and resection of the dura mater, 2 electrodes were each inserted into left and right hemispheres to target the dorsal HPC at [AP: −1.84 mm; ML: +/- 1.2 mm; DV: 1.4] via a stereotaxic frame (David Kopf Instruments, Tujunga, CA, USA). Real-time data display and an audio aid were used to facilitate the identification of proper recording locations while advancing electrodes until characteristic signals from stratum pyramidale and stratum radiatum were detected and recorded (He et al., 2020). A 1-hr multi-channel recording from bilateral sensorimotor cortex and dorsal HPC was collected from each mouse. Data were stored at a sampling rate of 32 kHz after band-pass filtering (0.1-9 kHz) with an input range of ± 3 mV (Digital Lynx SX, Neuralynx, USA). Data were down sampled to 1250 Hz for further analysis. A total of 23 mice were successfully recorded and subjected to data processing. The groups had the following counts: db/+ 200 d (n = 7), db/db 200 d (n = 6), db/+ 400 d (n = 5), db/db 400 d (n = 5). Mortality rate was about 15% due to reaction to urethane anesthesia.

### Electrophysiology data analysis

#### Spectral power analysis

Local field potentials from the pyramidal layer and stratum lacunosum moleculare (slm) of the HPC and the deep cortical layer of the sensorimotor cortex were used in our analysis. Brain waves were filtered from the LFPs according to the following frequency ranges: delta (0.1– 3 Hz), theta (4–7 Hz), gamma (30–58 Hz), and high-gamma (62–200 Hz) and signal power determined as previously described (He et al., 2020). A slowing score was calculated, defined as the ratio between low frequency (1-8 Hz) and high frequency oscillations (9-30 Hz), where higher values of the slowing score correspond to a shift in spectral power from high to low frequencies (Laptinskaya et al., 2020). Theta state in the brain was determined by calculating the ratio of theta/delta (T/D) Hilbert amplitudes in the slm layer of the HPC. The amplitude envelopes were smoothed with a Gaussian kernel (σ = 1 s, 8 s window), and the T/D was further smoothed with a second Gaussian kernel (σ = 10 s, 80 s window), to stabilize changes of state and reduce noise. The smoothed ratio was then split by a manual threshold set by visual assessment to define two states, high theta/delta (HT/D) and low theta/delta (LT/D) (Barth & Mody, 2011; Lockmann, Laplagne, Leao, & Tort, 2016; Wolansky, Clement, Peters, Palczak, & Dickson, 2006).

#### Estimation of the spectral exponent from the PSD background

Since changes in spectral power can be reflected by both periodic and aperiodic components, we also quantified the latter. Isolated from the aperiodic component of the signal as described previously (Donoghue et al., 2020), the spectral exponent (SE) measures the steepness of the decay of the power spectral density (PSD) background (Lanzone et al., 2022). PSD is assumed to decay according to the inverse power-law ∼1/*f*^α^, therefor we define the SE to be ꞵ = -α. The SE therefore is equivalent to the slope of the linear regression resulting from the log of the PSD (Colombo et al., 2019). We estimated the PSD using Welch’s method (2s window, 50% overlap) and SE calculated between the 1-40 Hz range.

#### SPW-Rs detection and characterization

SPW-Rs were identified from the pyramidal layer during LT/D periods (Ip et al., 2021). To isolate SPW-Rs, the LFP signal of the pyramidal layer was filtered (150–250 Hz), squared, and Z-scored. When the signal exceeded 4 standard deviations for a period longer than 0.05 msec, a SPW-R event was registered. When the signal subsequently dropped below 1 standard deviations, the event was considered to have ended. Multiple characteristics of SPW-Rs were calculated. <u>Duration</u> and inter-ripple interval (<u>IRI</u>) were calculated using the start and end timings of SPW-R events.

##### Pyramidal gamma LFP power during SPW-Rs

SPW-R-associated slow gamma signal power was calculated using the averaged z-scored power over the 30–50 Hz frequency band 0–100 msec after ripple detection. This was then averaged over all SPW-RS events.

#### Functional connectivity

Functional connectivity between the cortex and HPC areas was estimated using three methods; i.e. Phase locking index (PLI), coherence, and cross regional phase amplitude coupling (xPAC).

##### Phase locking Index (PLI)

An index of asymmetry of the distribution of phase differences between measured signals was calculated (Stam, Nolte, & Daffertshofer, 2007; Vinck, Oostenveld, van Wingerden, Battaglia, & Pennartz, 2011). PLI is a measure of the consistency in the distribution of instantaneous phase differences between two signals. If the phase differences between two time series are △ϕ(t_k)(k = 1 …N), PLI is defined as:

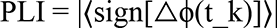

where 〈·〉 is the mean value operator. PLI values range from 0 - 1, where a value of 0 indicates no coupling or coupling with a phase difference centered around 0 mod pi, and a value of 1 reflects complete synchrony between two signals. PLI was computed for the following frequency bands: delta (0.1-3 Hz), theta (4-7 Hz), alpha (8-13 Hz), beta (13-30 Hz), gamma (30-58 Hz), and high gamma (62-200 Hz).

##### Coherence

Coherence measures the consistency of a phase relationship between two signals. Coherence was calculated pairwise between cortex, CA1 pyramidal, and CA1 slm layer of the HPC for each of the frequency bands. The coherence between signals x and y is defined as the square of the cross-spectrum of the channels divided by the product of the power spectra of the individual channels:

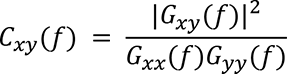

where Gxx and Gyy refer to power spectral density of channels x and y respectively, and Gxy refers to their cross-spectral density (Bendat & Piersol, 1986).

##### Cross-regional phase-amplitude coupling (xPAC)

xPAC was measured between the pyramidal layer and the cortex as described previously (He et al., 2020; Ip et al., 2021). To estimate xPAC we bandpass-filtered the LFP between 0.1 and 200 Hz, extracted the instantaneous phase from the pyramidal layer and instantaneous amplitude from the cortical layer using the Hilbert transform. A composite phase-amplitude time series then determined the amplitude distribution across phase. The modulation index (MI) was then calculated from the divergence of the amplitude distribution from a uniform distribution (Tort, Komorowski, Eichenbaum, & Kopell, 2010). MI was compared between groups by averaging the MI across a window of frequencies pertaining to canonical frequency bands. A data driven threshold was found using Otsu’s method (McIntyre et al., 2010) to determine the window of significant coupling.

### In vivo labeling by 5-bromo-2’-deoxyuridine-5’-monophosphate (BrdU)

To track cell proliferation, BrdU (Sigma-Aldrich, MO, USA) was administered intraperitoneally (i.p.; 50 mg/kg; two times per day for a period of 4 days) to the db/db and db/+ mice as described previously (C. Sun et al., 2013). To enhance the detection rate of colocalization between BrdU and cell type specific markers, the dose of BrdU was increased to 75 mg/kg and three times daily for a period of 4 days in the 200-day groups. Mice were euthanized 2-3 weeks after BrdU administration.

### Brain dissection and tissue processing

Prior to the brain dissection, adult mice were anesthetized with isoflurane (2-5% with 30-40% O2 and 60-70% nitrous oxide in gas), and intracardially perfused with 300 ml of 4% paraformaldehyde (PFA). The brains were extracted and post fixed in 4% PFA overnight at 4 °C. Following cryoprotection in 20% sucrose, 40 μm-thick coronal sections were cut on a freezing microtome and collected serially (Hong et al., 2007).

### Immunocytochemistry and immunofluorescence staining

Immunohistochemistry of dentate gyrus (DG) and subventricular zone (SVZ) regions was performed on 40-μm serial free-floating sections. To improve the efficiency of BrdU detection, sections were pretreated with 1 N HCl for 30 min and then neutralized with 0.1 M sodium borate buffer pH 8.5 for 10 min prior to incubation with primary antibody. Sections were blocked and permeabilized with blocking serum (0.3% Triton X-100, 2% BSA, and 1% donkey serum) for 30 minutes, followed by incubation in mouse anti-BrdU (1:400, Roche, Basal, Switzerland), goat anti-DCX (1:1000, C18, Santa Cruz Biotechnology, TX, USA), mouse anti-NeuN (1:1000, Millipore, MA, USA) and rabbit anti-GFAP (1:500, Dako, CA, USA) overnight at room temperature and then with donkey anti-mouse, anti-goat, or anti-rabbit secondary antibodies conjugated to Alexa 488 or 594 (1:400, Invitrogen, MA, USA) for 2 hours at room temperature according to the methods described previously (Z. Liu et al., 2007; C. Sun et al., 2013). For DAB staining, donkey anti-goat or anti-mouse biotinylated secondary antibodies were incubated for 2 hours at room temperature (1:1000, Jackson ImmunoResearch, West Grove, PA, USA), followed by incubation with VECTASTAIN Elite ABC HRP solution (Vector, CA, USA) for 1.5 hour at room temperature. The substrate response was detected with DAB (Sigma-Aldrich, MO, USA) until satisfactory brown staining was achieved. The sections were then dehydrated in ascending concentrations of alcohol, cleared in Citrisolv, mounted with permanent mounting medium as previously described (Fan, Liu, Weinstein, Fike, & Liu, 2007).

### Imaging and quantification

Fluorescence signals were detected by using the Zeiss Spinning Disk confocal image system (Zeiss, Thornwood, NY) using a sequential scanning model with step size of 1 µm for Alexa 488 and 594. Images were process by Zeiss ZEN software for orthogonal and maximal intensity projection. Montages were created by Adobe Photoshop (Adobe System, Mountain View, CA). DAB or cresyl violet (CV) signals were captured with a CCD camera attached to Zeiss Axio II microscope equipped with StereoInvestigator software (MicroBrightField, VT, USA).

Quantification of CV and DAB staining of DCX or BrdU in SVZ was performed as previously described (Parent, Vexler, Gong, Derugin, & Ferriero, 2002). Images were digitally captured under a 10X objective and imported into Image J (NIH) for analysis. The area of DCX or progenitor cell staining in dorsolateral SVZ in DAB and CV staining were outlined and calculated. The average cell count or area were averaged from 3 sections of SVZ and DG each per mouse at the same AP levels. DG cell counts were adjusted to represent the entire HPC by multiplying the total number of sections.

### Statistical analysis

We expressed data as mean ± standard deviation in all figures and performed two-way analysis of variance (ANOVA) to assess the main effects of age and T2DM, followed by Tukey’s post-hoc test to determine between-group differences in which the adjusted p values less than 0.05 were considered as significant.

## Results

### T2DM slowed neural rhythms in the HPC while age reduced T/D ratio

We assessed the slowing score, defined by the ratio of low frequencies (1-8 Hz) over high frequency oscillations (9-30 Hz) within the cortex or HPC. We found that T2DM was associated with significantly increased slowing score in both pyramidal (p < 0.05) and slm (p < 0.001) layers of the HPC (Fig. 1A). With respect to the effect on individual frequency band, T2DM had a tendency to lower signal power of higher frequency oscillations (Supplemental Fig. 1). Regarding brain state, age significantly decreased the ratio of HT/D to LT/D (p < 0.05), indicating that age reduced time spent in HT/D (Fig. 1B).

**Figure 1.**
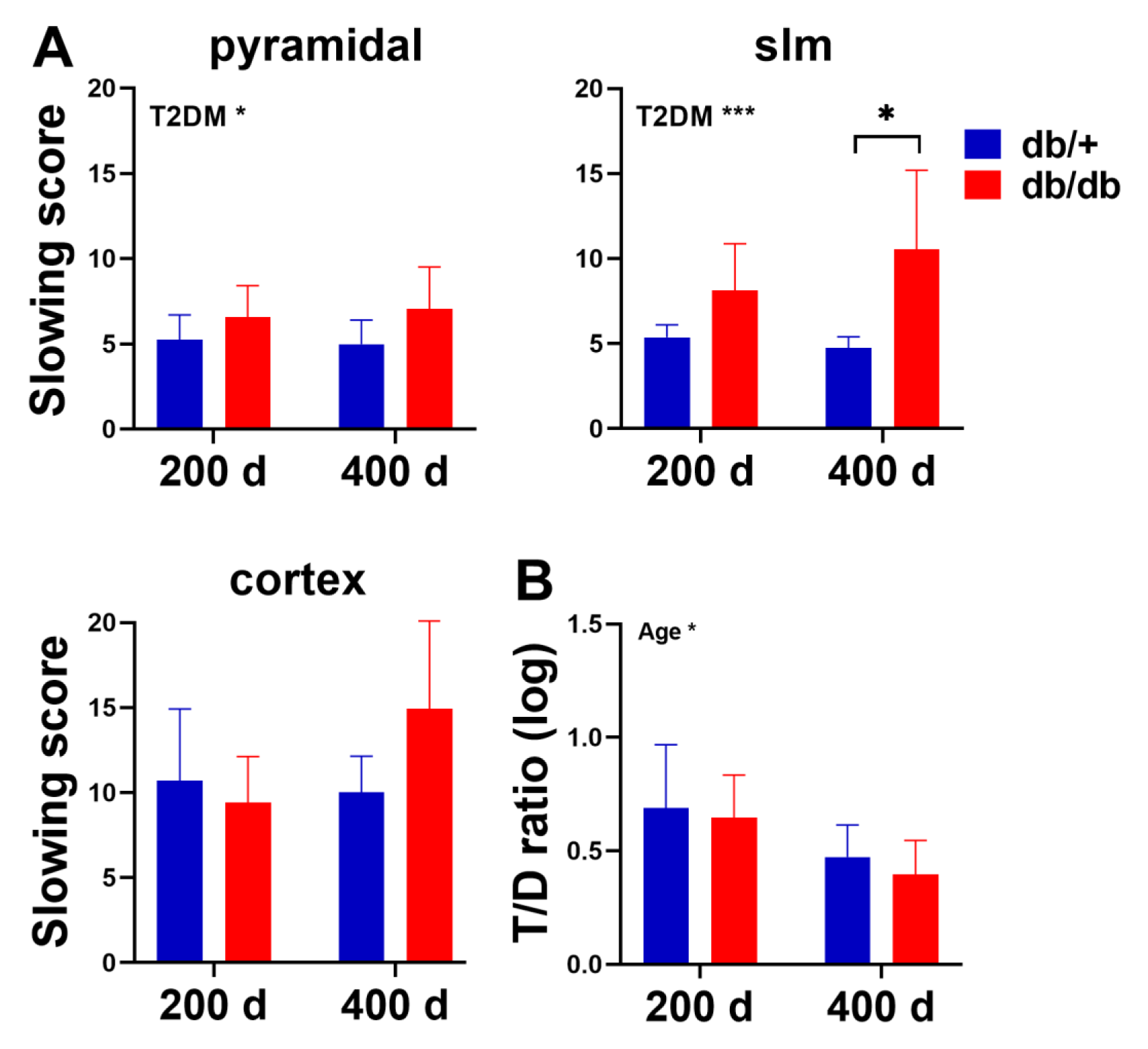
T2DM increased slowing score in the hippocampus (HPC) and age altered brain state. A, Diabetic mice shifted the power towards lower frequency oscillations in comparison to the control mice in the pyramidal and slm layers of the HPC. B, Age altered brain state by reducing theta/delta (T/D) ratio. Two-way ANOVA test followed by Tukey’s post-hoc test. 200 d: 200 days old, 400 d: 400 days old. *p < 0.05, ***p < 0.001.

### The spectral exponent of the aperiodic signal

We next analyzed the power spectral density and aperiodic signals of the field potentials. T2DM exponentially reduced power with increasing frequency in the slm layer of the HPC, reflecting a diabetes-associated increase in spectral exponent due to a faster decay of high frequency signal power in the HPC, which was more prominent in the older group (Fig. 2).

**Figure 2.**
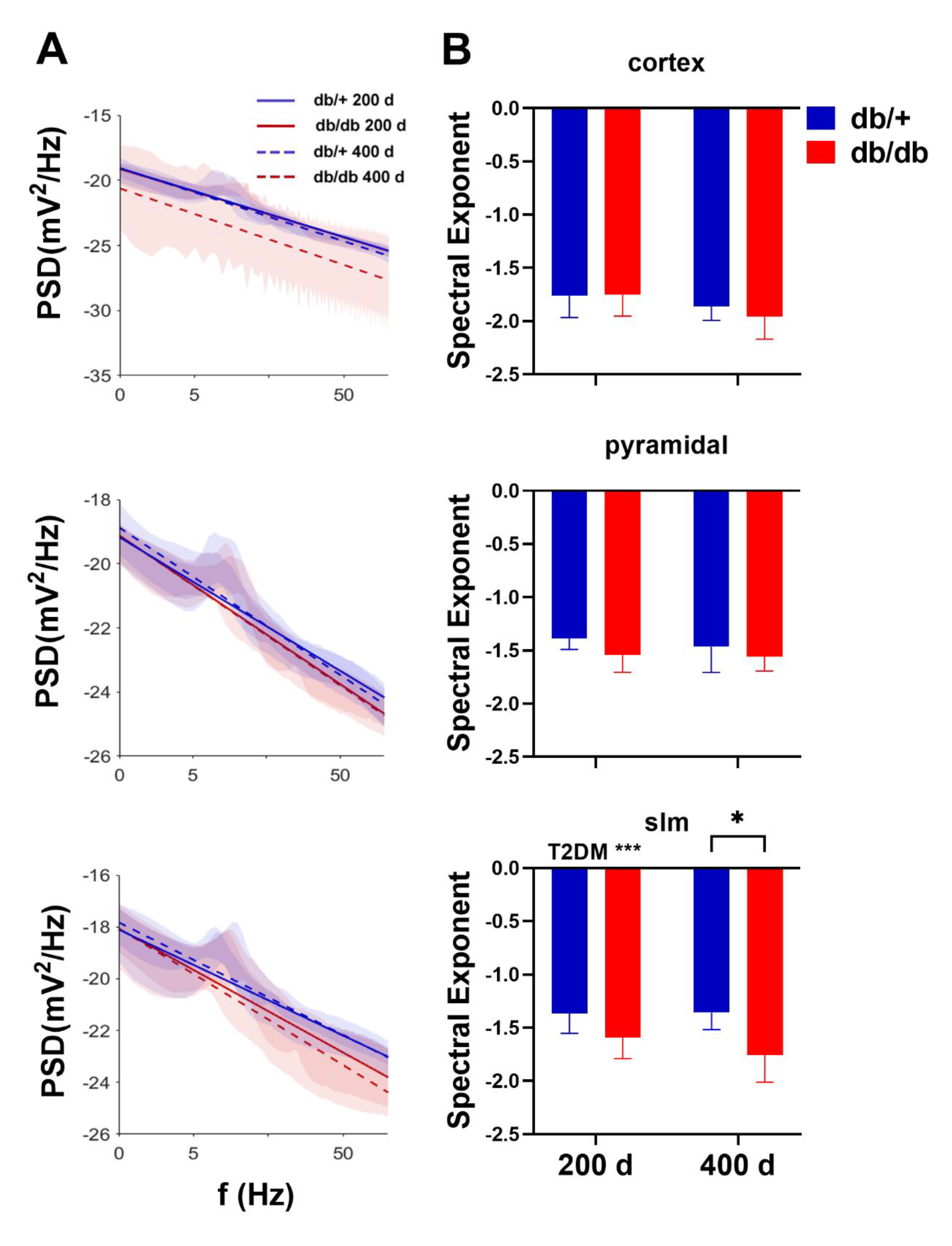
T2DM reduced spectral exponent in the slm layer of the HPC. A, Average power spectral density of recordings and best-fit trendline for each group. B, Comparison of the spectral exponent showing the older T2DM mice had the most reduced spectral exponent. Two-way ANOVA test followed by Tukey’s post-hoc test. 200 d: 200 days old, 400 d: 400 days old. *p < 0.05, ***p < 0.001.

### Age and T2DM reduced cortico-hippocampal coherence and phase synchrony

We determined functional connectivity between brain networks in the frequency domain by coherence and phase synchrony. By measuring the consistency of relative amplitude and phase between signals detected in two regions within a set of frequency band, we found that coherence significantly decreased as a function of age between cortex and CA1 in theta, alpha, beta, gamma (Two way ANOVA, age effect: p<0.05), and high gamma (age effect: p<0.01) frequency bands (Fig. 3A). Meanwhile, coherence reduced as a function of T2DM in alpha (T2DM effect: p < 0.05), beta (T2DM effect: p < 0.001), and high gamma (T2DM effect: p < 0.05) frequency bands between cortex and slm layer (Fig. 3B).

**Figure 3.**
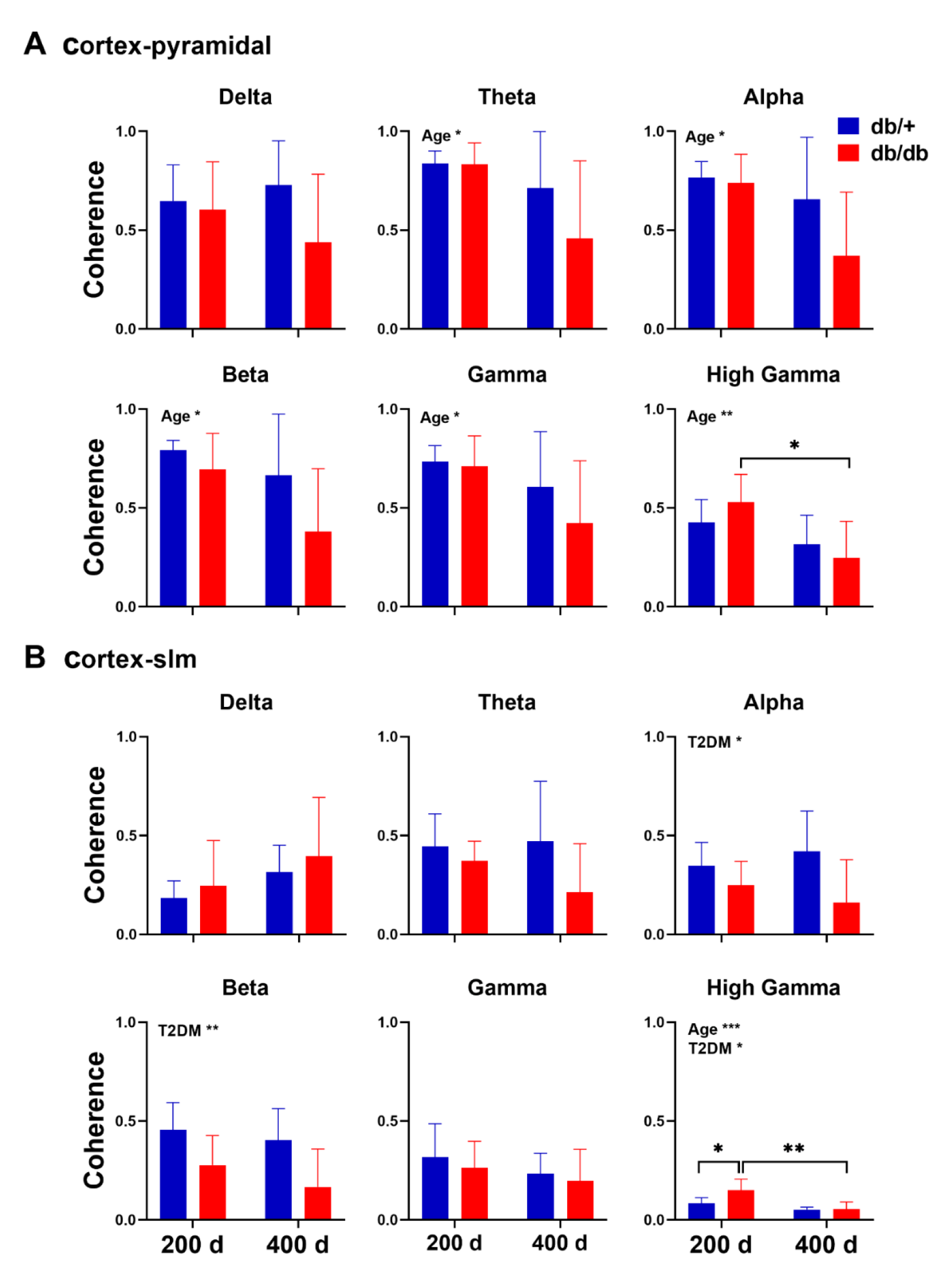
Age and T2DM reduced coherence at a number of frequency bands between cortex and HPC. A, Cortex-pyramidal coherence: Age decreased coherency between the cortex and pyramidal layer in all but delta frequency. B, Cortex-slm coherence: T2DM effect in reducing coherency was observed in alpha, beta, and high gamma frequency bands. Two-way ANOVA test followed by Tukey’s post-hoc test. 200 d: 200 days old, 400 d: 400 days old. *p < 0.05, **p < 0.01, ***p < 0.001.

By measuring how stable the phase difference varies over a period of time between two regions independent of the amplitude of oscillations, we found that the phase synchrony represented as PLI decreased as a function of age in theta, alpha, beta, gamma, and high gamma frequencies between cortex and CA1 (age effect: p<0.05) in a consistent manner as in coherence, in addition to the T2DM associated decrease in delta frequency (T2DM effect: p<0.05) (Fig. 4A). PLI also decreased as a function of T2DM in beta and gamma (T2DM effect: p<0.05) frequencies between cortex and slm layer (Fig. 4B). Our data suggest that both age and T2DM decreased functional connectivity between cortex and HPC.

**Figure 4.**
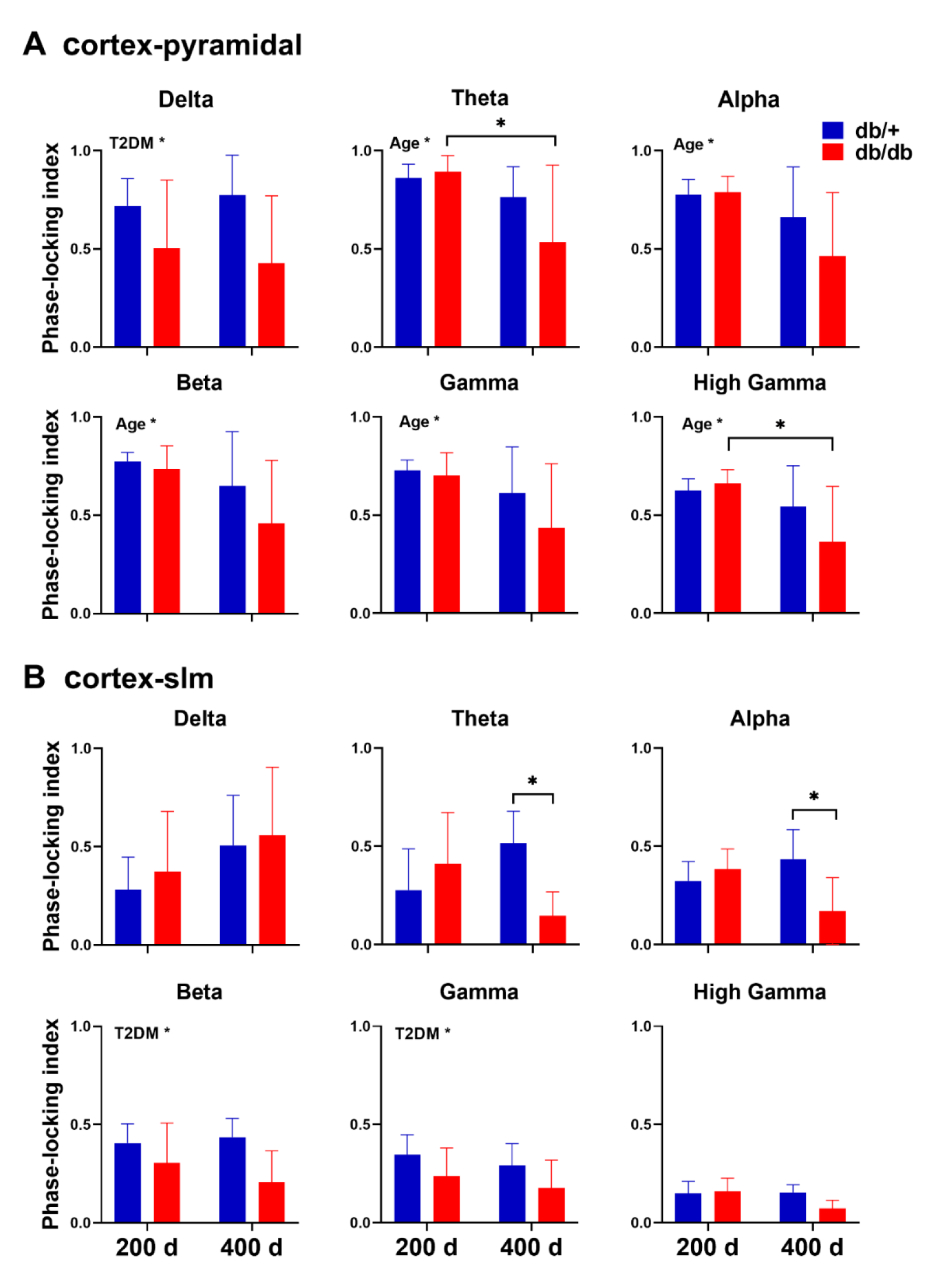
PLI calculation for individual frequency bands between A, cortex and pyramidal and B, cortex and slm layers. Age significantly decrease PLI between the cortex and pyramidal layer in all frequency bands except of delta frequency. T2DM decreased PLI between cortex and pyramidal layer in delta and between cortex and slm layer in theta, beta, and gamma bands. Two-way ANOVA test followed by Tukey’s post-hoc test. 200 d: 200 days old, 400 d: 400 days old. *p < 0.05.

### Both age and T2DM showed dynamic effects on cross regional phase-amplitude modulation between cortex and HPC

A third approach to assess functional connectivity is via cross regional phase-amplitude relationship in different frequency bands between cortex and pyramidal or slm layers of the HPC using phase-amplitude coupling analysis. We found that both age and T2DM increased delta-gamma phase-amplitude coupling between the HPC and cortex as shown by the MI, while T2DM decreased theta-gamma coupling (Fig. 5). Our data suggest that T2DM led to disrupted communication between cortico-hippocampal circuits.

**Figure 5.**
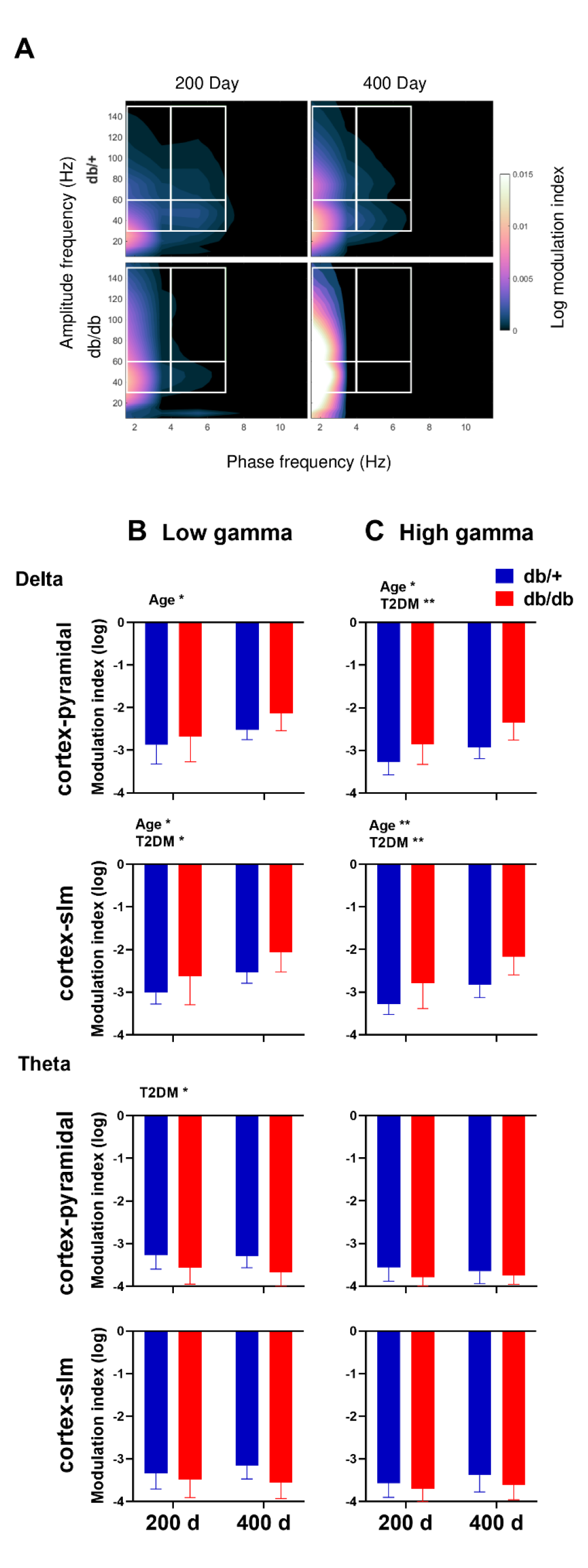
Cross regional phase-amplitude coupling between cortex and pyramidal or slm layer of the HPC. A, Comodulograms of cortex-pyramidal areas demonstrating delta-high gamma (upper left), theta-high gamma (upper right), delta-low gamma (lower left), and theta low-gamma (lower right) coupling area marked by white boxes. B, Quantified average log modulation index (MI) within areas of interest. Age or T2DM increased MI between delta and gamma, while T2DM reduced MI between theta and gamma. Two-way ANOVA test followed by Tukey’s post-hoc test. 200 d: 200 days old, 400 d: 400 days old. *p < 0.05, **p < 0.01.

### Age and T2DM affected the duration of SPW-Rs and gamma power during SPW-Rs

We next examined how age or T2DM affected the properties and emergence of SPW-R, a hippocampal specific oscillation resulting from the dynamical interaction between pyramidal cells and GABAergic interneurons within the local hippocampal circuits. We first examined the amount of network activation in CA1 during SPW-Rs by measuring gamma signal power (Fig. 6B). We found that both age and T2DM increased gamma power during SPW-Rs firing (Fig. 6B). We then found that both age and T2DM prolonged ripple duration (p < 0.0001; Fig. 6C). Specifically, older db/db mice had reduced IRI (Fig. 6D) compared to younger ones, likely due to increased ripple duration. The longer ripples and increased gamma power suggest that age and T2DM likely increased the excitability of the HPC.

**Figure 6.**
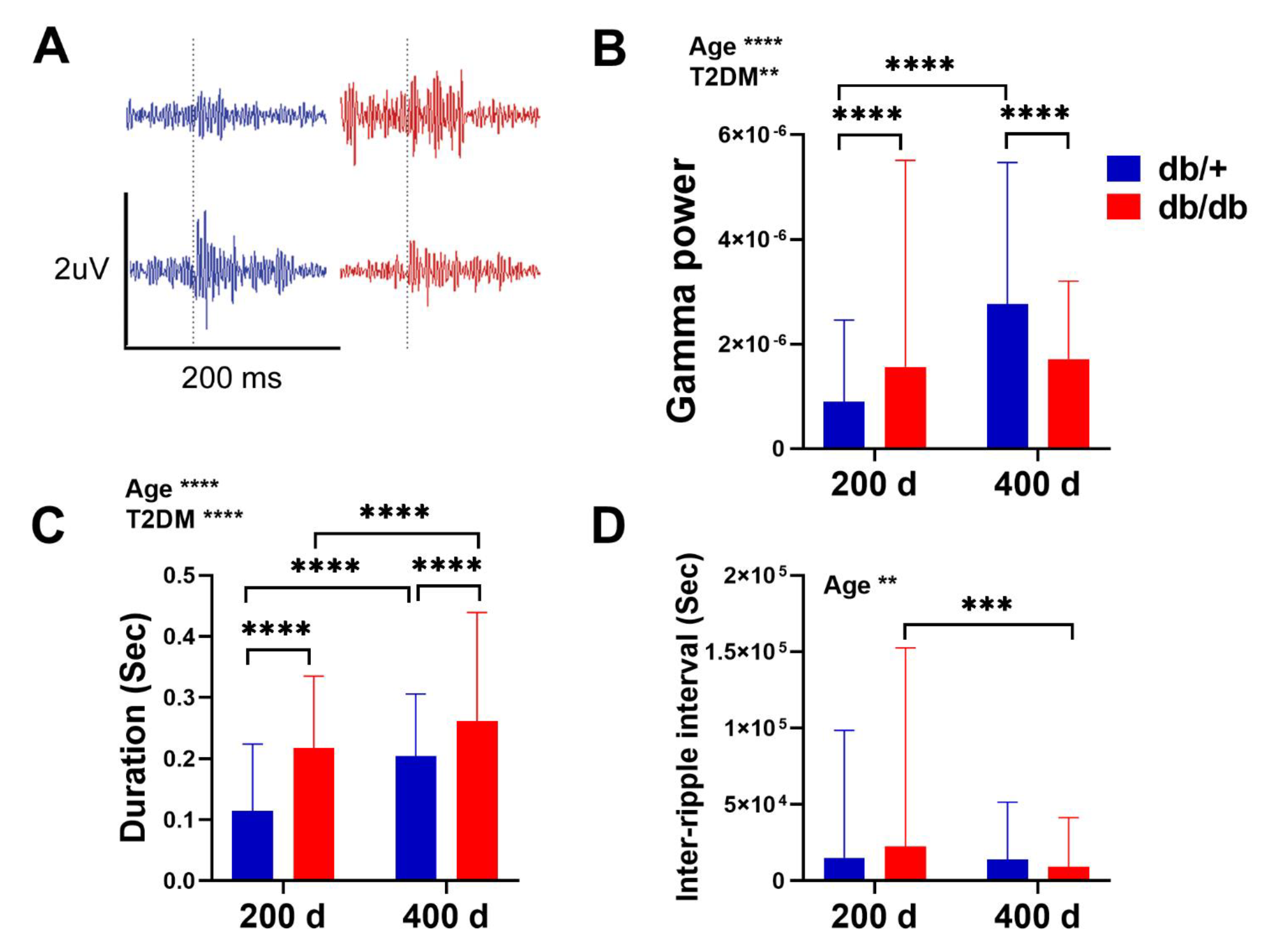
Characteristics of sharp wave associate ripples (SPW-Rs). A, Average ripple waveform in 200 msec clips. B, Comparison of gamma signal power during SPW-Rs. C, Comparison of duration of SPW-Rs. Age and T2DM increased ripple duration and gamma power during ripples. D, Comparison of inter-ripple intervals (IRI). Age significantly reduced IRI and this reduction was especially prominent in T2DM mice. Two-way ANOVA test followed by Tukey’s post-hoc test. 200 d: 200 days old, 400 d: 400 days old. An average of 220 ripples were analyzed per group. **p < 0.01, ***p < 0.001, ****p < 0.0001.

### Both age and T2DM decreased neurogenesis in the HPC

To determine how T2DM interacts with age to affect neural stem cell activity, we quantified the levels of neurogenesis in the SGZ of the dentate gyrus and the subventricular zone (SVZ). We found that both age and T2DM were associated with significant decreases in cell proliferation (p < 0.0001; Fig. 7A) and the total number of DCX(+)-neuroblasts in the SGZ (p < 0.0001; Fig. 7A). Newborn cells from 200- day control and diabetic mice developed into neurons and astroglia in the dentate gyrus 2-4 weeks after the division of progenitor cells (Fig 7B, C). There was also a significant age-associated reduction in neurogenesis in the SVZ (p < 0.01; Fig. 8).

**Figure 7.**
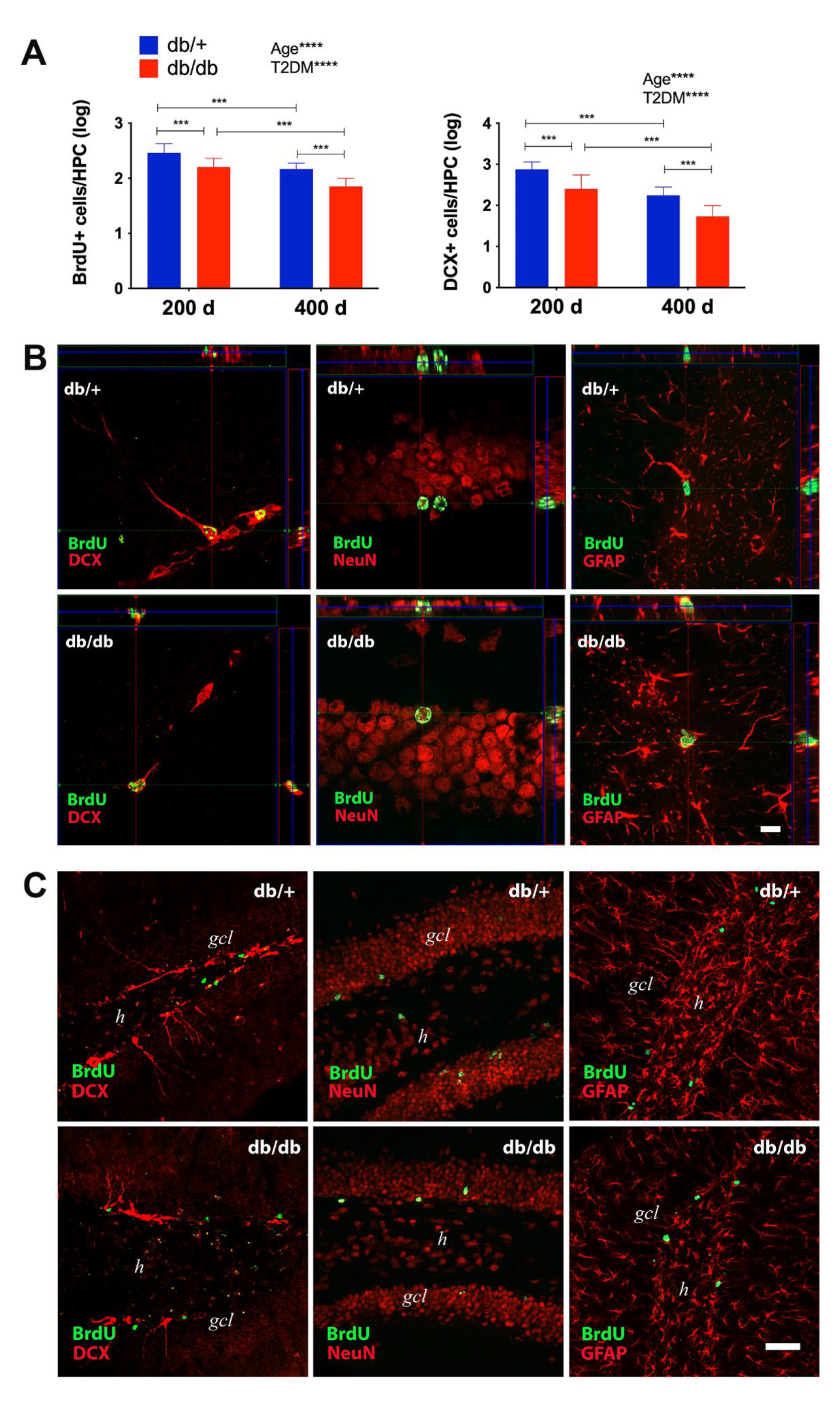
Age and diabetes reduced cell proliferation and immature neurons in the dentate gyrus subgranular zone (SGZ). The numbers of proliferating neural progenitor cells and neuroblasts in the SGZ of the HPC were quantified using (A) double cortin (DCX) and BrdU staining. Two-way ANOVA test followed by Tukey’s post-hoc test showed significant effect of age and T2DM on hippocampal neurogenesis. 200 d: 200 days old, 400 d: 400 days old. ***p < 0.001, and ****p < 0.0001. B, Representative double immunofluorescent staining of the dentate gyrus in lower magnification (20X) view from 200-day old db/+ (upper panel) and db/db (lower panel) for BrdU (green) and DCX, NeuN or GFAP (red). gcl: granule cell layer. h: hilus. Scale bar, 50 µm. C, Orthogonal reconstructions of confocal microscope merged images from db/+ (upper panel) and db/db (lower panel) with BrdU as green and cell markers (DCX, NeuN or GFAP) as red as viewed in the x–z (top) and y–z (right) planes. Scale bar, 10 µm.

**Figure 8.**
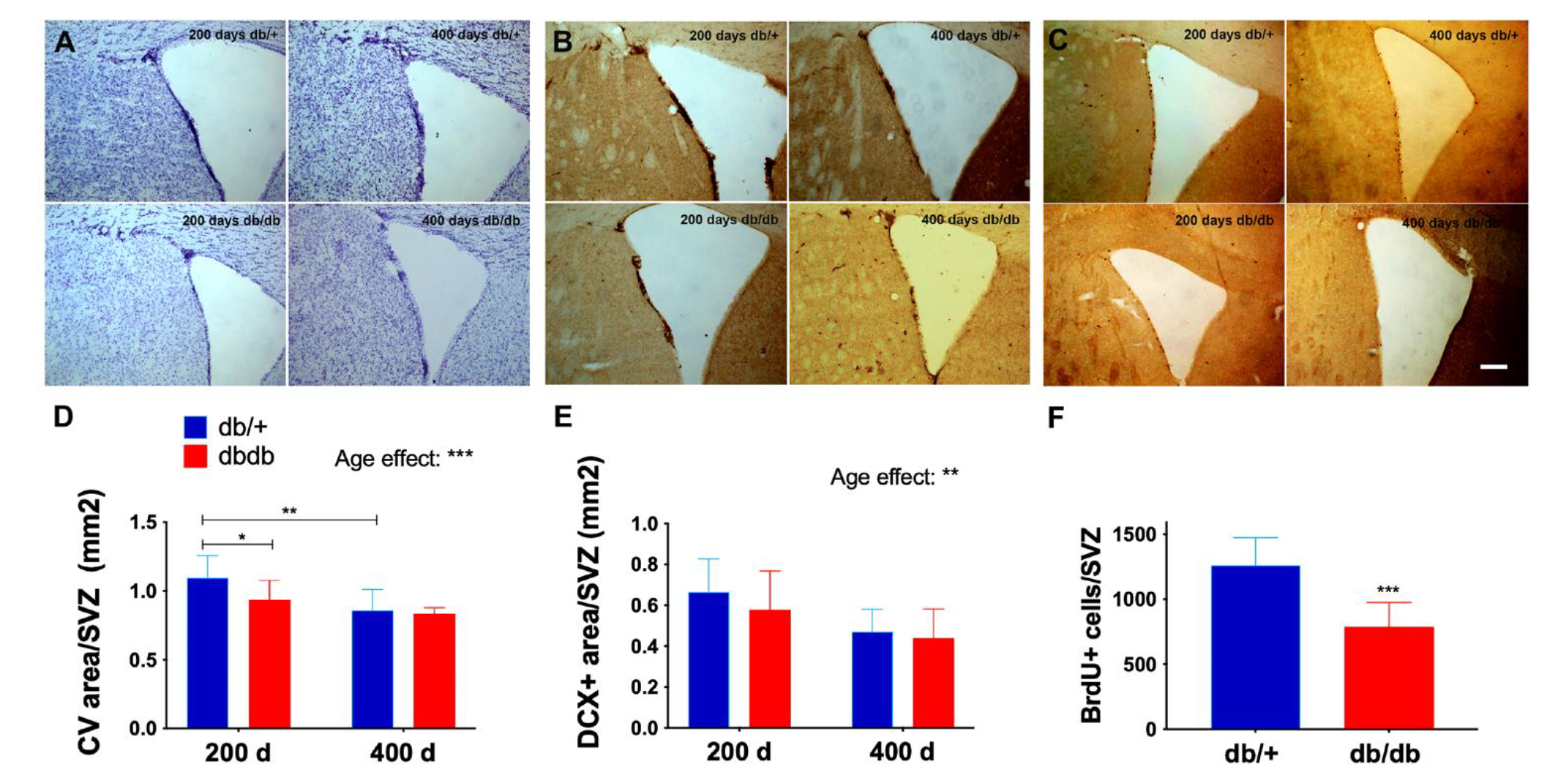
Age reduced progenitor cell activity in the subventricular zone (SVZ). The number of proliferating neural progenitor cells and neuroblasts in the SVZ was quantified using Cresyl violet (CV) staining (A, D) DCX staining (B, E) and BrdU staining (C, F). Representative images (A-C) from each group showed the regions in the SVZ and lateral ventricle where neuroprogenitor cells and neuroblasts reside. Scale bar, 100 µm. BrdU counts were not shown for the 400 d groups due to very low numbers were found (F). There was an overall effect of age on neuroprogenitor cells and immature neurons, while T2DM reduced neuroprogenitor cells and cell proliferation in the 200 d group. Two-way ANOVA test followed by Tukey’s post-hoc test. * p < 0.05, ** p < 0.01, and *** p < 0.001. 200 d: 200 days old, 400 d: 400 days old.

## Discussion

While systematic changes in neural oscillations are known to occur under normal and pathological aging (Giovanni et al., 2017; Marshall & Cooper, 2017; Neto, Biessmann, Aurlien, Nordby, & Eichele, 2016; Rossini, Rossi, Babiloni, & Polich, 2007; Vlahou, Thurm, Kolassa, & Schlee, 2014; Voytek & Knight, 2015), how cognitive aging risk factors alter or accelerate these natural progressions is unclear. Here, we investigated the effect of T2DM and aging on neural oscillations in the HPC and neural synchrony between brain regions. We found that age strongly reduced theta brain state, while T2DM significantly increased slowing score and reduced the spectral exponent of the aperiodic signal in the HPC. Age and T2DM both increased SPW-Rs duration and gamma power during ripple firing. With respect to neural synchrony, age and T2DM reduced PLI and coherence between cortex and HPC in various frequencies. Most importantly, both age and T2DM increased delta-gamma coupling while T2DM reduced theta-gamma coupling between HPC and cortex. Although T2DM is widely accepted as an accelerated aging with known effect on cortical atrophy, we report here for the first time that the electrophysiological manifestation of T2DM is more prominent in the HPC as reflected by the increased SPW-Rs and slowing score as well as reduced neural synchrony between cortex and HPC.

Reduced amplitude and peak frequency of the alpha-band was often reported during normal cognitive aging in humans (Knyazeva, Barzegaran, Vildavski, & Demonet, 2018; Marshall & Cooper, 2017; Mierau, Klimesch, & Lefebvre, 2017), whereas our study in mice showed a consistent age effect on diminishing signal power in theta and alpha bands in the cortex. Compared to age effect, T2DM significantly increased slowing score and decreased signal power in the beta frequency range in the HPC. In addition to changes often reported in the periodic component, recent work has suggested that the aperiodic component of LFP signals contains physiologically relevant information that changes with age and cognitive functions (Hill, Clark, Bigelow, Lum, & Enticott, 2022). Our data in T2DM-associated steepening in the spectral exponent of the aperiodic signal implies a tilted balance of synaptic excitation and inhibition (Gao, Peterson, & Voytek, 2017; Guo et al., 2018). Apart from weakening oscillation power, we found that age also altered brain state by proportionally prolonging the LT/D periods, and in turn shortening HT/D periods. The balance and duration of brain states has been shown to be essential to proper memory function (Buzsáki, 2002), and thus an altered brain state may be an indicator of disrupted learning, memory, and cognitive functions.

As one of the electrophysiological proxies of functional connectivity between brain regions, PLI measures neuronal synchrony between two recorded signals. A reduction in PLI globally across the neocortex has been reported in T2DM (Zeng et al., 2015) and mild cognitive impairment (Kuang et al., 2022; Youssef et al., 2021). In our study we have also detected a decrease in PLI between cortex and HPC over various frequency bands as a function of T2DM or age, showing that the decrease in PLI extends to connections between the neocortex and the temporal lobe in diabetic or aged mice. Unlike PLI which measures the consistency of the instantaneous phase difference between two signals, coherence determines the ratio of the cross spectral density and the individual auto spectral densities albeit with a lower temporal specificity. Coherence has been shown to be altered by neurological disorders such as stroke (Cassidy et al., 2020), depression (Danish M Khan et al., 2022; D. M. Khan, Yahya, Kamel, & Faye, 2023), AD (Rodinskaia, Radinski, & Labuhn, 2022) and Down syndrome with AD (Musaeus, Salem, Kjaer, & Waldemar, 2021). Similar to the PLI, we detected a decrease in coherence due to age or diabetes. Theta oscillations in the HPC provide a temporal reference for gamma oscillations through theta-gamma coupling (Buzsaki, 2015). Consistent with recent work revealing that the HPC and cortical areas utilize xPAC to support memory encoding (Wang, Schmitt, Seger, Davila, & Lega, 2021), our data showed that theta-gamma xPAC was reduced between cortex and HPC by T2DM, similar to the effect of chronic ischemic stroke in rats (Ip et al., 2021). Our findings in T2DM associated reduction in functional connectivity in mice assessed by electrophysiology is in line with the fMRI findings in human T2DM patients (D. Liu et al., 2018; Q. Sun et al., 2018; Zhang et al., 2015; Zhou et al., 2010).

SPW-Rs are among the most synchronous spontaneous population patterns in the mammalian brain, and recent evidence suggest that these waves serve to reactivate neurons encoding episodic memories to promote memory consolidation and also contribute to the planning of future actions by generating ordered neuronal firing sequences (Buzsaki, 2015; Foster, 2017; Oliva, Fernández-Ruiz, de Oliveira, & Buzsáki, 2018). SPW-Rs may be phase-coupled with a power spectral peak in the slow gamma band originating from the CA3, which in turn determines information flow in the HPC-EC system (Kitanishi et al., 2015). Thus, disruption of SPW-Rs and/or gamma oscillation in the HPC-EC of experimental animals and humans causes severe memory impairment (Fernandez-Ruiz et al., 2019; Fernández-Ruiz et al., 2021; Hollnagel et al., 2019; Jadhav, Kemere, German, & Frank, 2012; Jones, Gillespie, Yoon, Frank, & Huang, 2019; Le Van Quyen et al., 2010). For the first time our study has revealed that age and T2DM increase the duration of hippocampal SPW-Rs and gamma power during SPW-Rs, while age reduce inter-ripple interval. The increased gamma power during SPW-Rs suggest that neurons firing during SPW-Rs become more excitable as a function of age or T2DM.

Our finding in age-and T2DM-associated reduction in hippocampal neurogenesis and changes in SPW-Rs characteristics is consistent with the physiological role of hippocampal neurogenesis in maintaining the balance of excitatory and inhibitory activity and proper cognitive function. Established evidence suggests that newborn neurons project monosynaptic inhibitory input onto granule cells, producing a feed-forward inhibition of CA3 neurons (Luna et al., 2019). This process is crucial in maintaining remote memory (Guo et al., 2018), but is weakened by age (Oh, Simkin, & Disterhoft, 2016), resulting in hyperexcitability of the CA3 auto-associative network that has been proposed to lead to memory rigidity during aging (Leal & Yassa, 2015; Wilson et al., 2006). Increased neurogenesis by Cdk4/cyclinD1 overexpression triggered an overall inhibitory effect on the trisynaptic hippocampal circuit and reversed age-associated CA3 hyperactivation, resulting in decreased occurrence, duration and increased interval of SPW-Rs (Berdugo-Vega et al., 2020). However, our findings might not seem intuitively compatible with an earlier study showing that the longer duration of ripples was found to be related to mnemonic demand and performance (Fernandez-Ruiz et al., 2019). A few factors might have contributed to the seemly discrepant findings between the two studies. First, the main difference is that the SPW-Rs detected in our study are not task dependent. Second, the average duration of SPW-Rs detected in our T2DM or older mice was more than 250 msec, which was significantly longer than the physiological ripples ranging 1-200 msec. Longer ripples likely reflect the pathological condition of the HPC and differ from physiological ripples in their involvement in memory function. In the condition of traumatic brain injury, the resulting deafferentation was reported to induce hyperexcitability of distal dendrites in the hippocampal pyramidal neurons (Cai et al., 2007). Consistent with this notion, longer duration of SPW-Rs were also observed in our ischemic stroke model (Ip et al., 2021), in which synaptic input from entorhinal cortex to the dentate gyrus was affected due to cortical injury (Sato et al., 2022; C. Sun et al., 2013).

Ample evidence supports a bidirectional relationship between diabetes mellitus (DM) and major depressive disorder (MDD) in humans (Chien, Wu, Lin, Chou, & Chou, 2012; Demakakos, Pierce, & Hardy, 2010; Kan et al., 2013; Khaledi, Haghighatdoost, Feizi, & Aminorroaya, 2019; Lloyd, Pambianco, & Orchard, 2010; Pan et al., 2010; Zhu et al., 2022). Shared abnormal neurophysiological features between patients with DM and depression are well documented including elevated power in delta and theta bands, and impaired response to task-oriented stimulation such as increased P300 latency in EEG (Baskaran, Milev, & McIntyre, 2013). Interestingly, overlapping pathology such as the pattern of volumetric abnormality and neurocognitive deficits was also found between diabetic patients and those with depressive disorder (McIntyre et al., 2010). Consistent with clinical evidence, depression-like behavior was detected in preclinical T2DM model db/db mice by forced swim test accompanied by thigmotaxis behavior and hypo-locomotion at a relatively young age of 10-11 weeks (Sharma, Elased, Garrett, & Lucot, 2010). The role of impaired leptin production or signaling in depression was supported by a study in which treatment of diabetic mice with leptin reversed the depressive-like behavior in the tail suspension test (Hirano, Miyata, & Kamei, 2007). However, the pharmacology and pathophysiology of leptin signaling defect in causing depression is not well understood. It is possible that leptin defect could cause depression by modulating the firing and downstream signaling of monoaminergic neurons in the forebrain. This is in line with the evidence that leptin receptor is expressed in the serotonergic raphe nuclei (Finn, Cunningham, Rickard, Clifton, & Steiner, 2001), and the leptin deficient ob/ob mice have reduced serotonin transporter expression in raphe nuclei (Collin, Hakansson-Ovesjo, Misane, Ogren, & Meister, 2000). Leptin also increased the production of forebrain 5-hydroxyindoleacetic acid, a breakdown product of serotonin (Calapai et al., 1999). In addition, systemic leptin treatment reversed the hedonic-like deficit induced by chronic stress and produced an antidepressant-like effect in the forced swim tests in rats (Lu, Kim, Frazer, & Zhang, 2006). Interestingly, the authors found that the targeted brain regions of leptin intervention are in the HPC and amygdala including the dentate gyrus as mapped by Fos expression (Lu et al., 2006).

Several limitations are noted in our study. Limited recording sites is one major weakness of our electrophysiology study, which precludes an in-depth analysis of the disrupted functional networks by age and T2DM. Although the linear array used in our study reveals superior hippocampal rhythms compared to conventional EEG, lacking globally distributed recording sites does limit our assessment of functional connectivity to only restricted cortical and hippocampal networks. Besides, without task or event-related recording, data collected under urethane anesthesia have limited implication in physiological cognitive function. The other main limitation of our study is that we did not establish a causal relationship between reduced neurogenesis and perturbed electrophysiology in our T2DM model, which hinges upon future development of therapy to enhance or restore neurogenesis in the db/db mice.

## Author Contributions

G.R. performed recordings, data analysis and contributed to the writing of the manuscript. Z.I. carried out data analysis and contributed to writing of the manuscript. S.Z. performed data analysis validation. H.Z. contributed to neurogenesis analysis. Y.S. assisted in electrophysiology recording. A.Y. contributed to analysis methods and validated the analysis. J.L. conceived the study design, planning and supervision of the experiments, interpretation of results and writing of the manuscript. All authors have read and agreed to the published version of the manuscript.

## Institutional Review Board Statement

The study was conducted in accordance with the Guide for Care and Use of Laboratory Animals issued by the National Institutes of Health and approved by San Francisco Veterans Affairs Medical Center Institutional Animal Care and Use Committee (protocol number 20-015, approved 8-10-2020).

## Abbreviated title

Age and diabetes affect hippocampal oscillations and neurogenesis

## Supplemental Material

### Effect of age and type 2 diabetes on the signal power of the oscillation frequencies

To gain a deeper insight into the T2DM associated slowing score, we determined the individual signal power of each oscillation frequency within the deep layers (layers 4-6) of sensorimotor cortex, CA1 field pyramidal layer, and slm layer in the hippocampus. Following two-way ANOVA, we found that age significantly reduced signal power of theta (p < 0.01) and alpha (p < 0.05) in the cortex. Although T2DM had a tendency to lower signal power of higher frequency oscillations, significant diabetes effect was only found in beta power in the pyramidal layer (p < 0.05) (Supple Fig. 1).

**Supple Figure 1.**
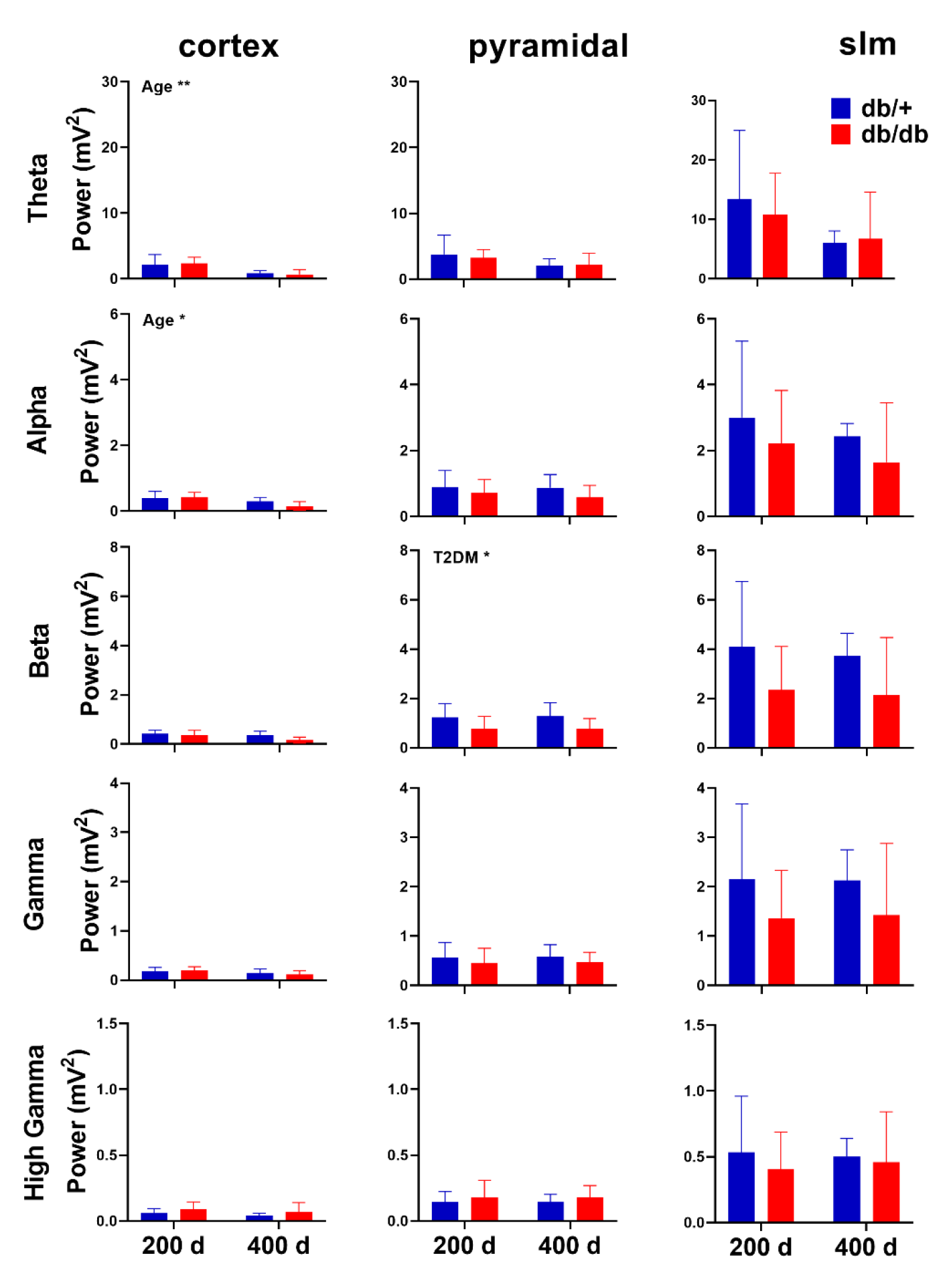
The effect of age and T2DM on signal power in the cortex and hippocampal layers. The main effect of age was observed in cortex for theta and alpha bands while the main effect of T2DM was limited to the beta frequency in the pyramidal layer of the hippocampus. Two-way ANOVA followed by Tukey’s post-hoc test. 200 d Tukey’s: 200 days old, 400 d: 400 days old. *p < 0.05, **p < 0.01.

